# ApicoTFdb: The comprehensive web repository of apicomplexan transcription factors and regulators

**DOI:** 10.1101/530006

**Authors:** Rahila Sardar, Abhinav Kaushik, Rajan Pandey, Asif Mohmmed, Shakir Ali, Dinesh Gupta

## Abstract

Despite significant progress in apicomplexans genome sequencing and genomics, the current list of experimentally validated TFs in these genomes is incomplete and mainly consists of AP2 family of proteins, with only a limited number of non-AP2 family TFs and TAFs. We have performed systematic bioinformatics aided prediction of TFs and TAFs in apicomplexan genomes, and developed ApicoTFdb database which consists of experimentally validated as well as computationally predicted TFs and TAFs in 14 apicomplexan species. The predicted TFs are manually curated to complement the existing annotations. The current version of the database includes 1310 TFs, out of which 833 are novel and computationally predicted TFs, representing 22 distinct families across 14 apicomplexan species. The predictions include TFs of TUB, NAC, BSD, CCAAT, HTH, Cupin/Jumonji, winged-helix, and FHA family proteins, not reported earlier in the genomes.

Apart from TFs, ApicoTFdb also classifies TAFs into three main subclasses-TRs, CRRs and RNARs, representing 3047 TAFs in 14 apicomplexan species are analyzed in this study. The database is equipped with a set of useful tools for comparative analysis of a user-defined list of the proteins. ApicoTFdb will be useful to the researchers interested in less-studied gene regulatory mechanisms mediating the complex life cycle of the apicomplexan parasites. The database will aid the discovery of novel drug targets to much needed combat the growing drug resistance in the parasites.

## Introduction

Transcription regulation is a key process that facilitates the cellular responses to different environmental conditions. The underlying transcriptional machinery of regulation is more complex in eukaryotes as compared to that in prokaryotes due to involvement of a diverse set of transcriptional enzymes and proteins acting as regulators. These regulators consist of site-specific Transcription Factors (TFs) as well as general TFs (TBP, TFIIB, TFIIE and MBF [1]) and specific RNA polymerases subunits [2]. In general, eukaryotic genomes contain a large number of TFs, classified on the basis of more that 90 kinds of conserved DNA binding domains (DBDs) [3]. The TFs can bind to specific DNA sequences upstream of promoter regions, controlling the rate of transcription and, thus transfer of genetic information [4, 5]. Here, the sequence diversity among DBDs also ensures precise regulation of various cellular processes in response to external and internal perturbations [6]. In fact, even in well-annotated organisms, numerous TFs have obscure DNA binding sequences which can still direct complex transcription regulation [7]. In addition, the transcription co-factors such as Chromatin Remodeling Factor, also control the direction of gene regulation by assisting general TFs [8]. Moreover, notwithstanding for the best-examined classes of DBDs, due to the diversity in protein as well as within the recognition sequences, the precise prediction of the regulators remains a challenging task [9].

This is especially important for recently sequenced or genomes with several annotated proteins with unassigned functions, for example *Plasmodium*, other apicomplexans like *Eimeria, Theileria* and *Cryptosporidium* genomes. However, despite the need, the number of annotated TFs in these apicomplexans genomes is exceptionally limited as compared to the model organisms like *H. sapiens, M. musculus*, and *A. thaliana* [10, 11]. In general, identification of TFs is either based on the experimental findings, for instance chip-seq, protein binding microarrays (PBM), or computational analysis by exploiting traditional sequence-similarity based search, e.g. BLAST and HMMER [12]. Herein, the computational methods compares the putative TF sequence with known DBDs as a reference for TF identification [13]. However, several TFs share a low sequence similarity with known DBDs [12], making their identification and characterization a daunting task, using traditional methods alone.

Till date, a number of TF databases have been developed-AnimalTFDB for animals [14], PlantTFDb for plants [15], FlyTF for fruit fly [16], TFCat [17] and TCOfDb [18] for human and mouse, however, there is no report of any database dedicated to apicomplexan specific TFs or their classified regulators. Using *in-silico* approach, Vaquero *et. al*. identified 202 transcription associated proteins in *P. falciparum* and classified them into general TFs, stage-specific TFs, and chromatin-related proteins [19]. However, only a limited number of TFs, mainly belonging to AP2 family, are experimentally validated in the parasite [10]. In virtue of their essential role in guiding the key cellular processes in the parasite’s life cycle involving multiple hosts, identification and characterization of novel TFs may provide deeper understanding of gene regulation in the parasite which may lead to identification of new drug targets. Therefore, in the present study, we performed an integrative *in silico* proteome analysis of 14 apicomplexan species in order to identify TFs and TAFs, based on conserved DBD analysis, interpro domain information [20] and gene ontology analysis [21]. We are able to identify and report several new TFs and TAFs belonging to diverse protein families among different apicomplexan species, including *P. falciparum*. Using this information, we developed ApicoTFdb-a novel web-based repository for hosting the classified list of apicomplexans regulators and information related to their domain architecture, molecular function(s), biological pathway(s), interlogs and dynamically linked to related literature. With this database, we highlight several putative regulators that otherwise remain obscure with existing resources.

We believe that the presented database would be extremely useful for the scientific community interested in deducing the regulatory molecules and their mechanism that governs the complex life cycle of any of these 14 different parasites. ApicoTFdb is freely accessible at http://bioinfo.icgeb.res.in/PtDB/.

## Materials and methodology

The protein sequences for TF identification across 14 apicomplexans species (see Table 1) were retrieved from PlasmoDB (version 38) [22], ToxoDB (version 38) [23], PiroplasmaDB (version 38) [24], and CryptoDB (version 38) [25]. We also obtained the sequences representing different classes of DBDs from AnimalTFDB (version 2.0), DBD (Release 2.0) and FootprintDb (as on 14/06/2018) [26]. These DBD sequences were used to create reference TF-HMM profiles with *hmm-search* program (HMMER version 3.1) [27]. To determine the set of DBD enriched protein sequences in the selected proteomes, we mapped each protein sequence to the reference HMM profiles (e-value < 0.001). Independently, we also searched conserved domains in each of the selected protein sequence using InterProScan5 (version 5.31) [20]. The results obtained from both of the independent methods were manually compared before assigning function to a given protein as putative TFs/TAFs (details in next sections). The additional annotation for the assigned TFs/TAFs, such as sequence length, gene and protein sequence, isoelectric point, molecular weight, previous IDs, and UniProt were retrieved from EupathDB. For each of the putative regulator, GO and biological pathway information were retrieved from AMIGO [28] and KEGG database [29]. The PPI information was retrieved from STRING databases (version 10.5) [30]. Nuclear Localization Signals were predicted using NucPred [31], and CELLO2GO was used for subcellular localization prediction [32]. PATS server was used to predict apicoplast-specific TFs [33].

**Table 1.** Updated transcription factors annotation in ApicoTFdb

| Species | Genome Size (in Mb ) | Total number of proteins (PlasmoDB ver.38) | PlasmoDB Text Search | Number of validated and predicted TFs in published databases |  |  |  | ApicoTFdb |  |  |  |
| --- | --- | --- | --- | --- | --- | --- | --- | --- | --- | --- | --- |
|  |  |  |  | DBD* |  | CisBP* |  |  |  |  |  |
|  |  |  |  | TF count | TF family | TF count | TF family | TF count | TF family | Experimentally verified | Unannotated proteins |
| *P. falciparum* | 23.3 | 5635 | 548 | 18 | 7 | 44 | 10 | 107 | 19 | 26 | 11 |
| *P. berghei* | 18.5 | 5089 | 418 | 19 | 12 | 46 | 11 | 99 | 20 | 13 | 15 |
| *P. chabaudi* | 18.8 | 5282 | 372 | 19 | 7 | 46 | 11 | 99 | 20 | 0 | 16 |
| *P. knowlesi* | 24.3 | 5261 | 386 | 14 | 8 | 46 | 10 | 86 | 18 | 12 | 9 |
| *P. vivax* | 28.8 | 5390 | 372 | 15 | 7 | 49 | 8 | 104 | 19 | 12 | 10 |
| *P. yoelii* | 22.4 | 7774 | 360 | 15 | 6 | 46 | 9 | 83 | 16 | 10 | 10 |
| *T. gondii*ME49 | 64.5 | 8920 | 443 | N.A* | N.A* | 81 | 14 | 146 | 18 | 1 | 12 |
| *T. gondii* p89 | 64.1 | 9874 | 486 | N.A* | N.A* | N.A* | N.A* | 145 | 18 | 0 | 4 |
| *B. bovis* | 8.1 | 3721 | 126 | N.A* | N.A* | N.A* | N.A* | 33 | 6 | 0 | 11 |
| *C. parvum* | 9.1 | 4020 | 248 | 30 | 11 | 35 | 8 | 77 | 13 | 1 | 5 |
| *C. cayetanensis* | 44 | 7592 | 332 | N.A* | N.A* | N.A* | N.A* | 67 | 13 | 0 | 35 |
| *N. caninum* | 57.4 | 7266 | 361 | N.A* | N.A* | 84 | 11 | 107 | 15 | 2 | 42 |
| *E. maxima* | 49.9 | 6249 | 241 | N.A* | N.A* | N.A* | N.A* | 74 | 18 | 0 | 46 |
| *E. tenella* | 51.8 | 8634 | 307 | N.A* | N.A* | N.A* | N.A* | 101 | 13 | 0 | 53 |
| *DBD- Transcription factor prediction database (Release 2.0)<br>*CisBP- Catalog of Inferred Sequence Binding Preferences (Last updated: Apr 5th, 2015 Database Build 1.02)<br>*N.A - No information available |  |  |  |  |  |  |  |  |  |  |  |

### TFfamily assignment

In order to classify a given protein sequence into a TF-family, we exploited its DBD profiles using the methods mentioned before. For a given protein, we independently obtained its domain information predicted with interproscan5 and GO based biological function, if available. We performed careful manual curation for each sequence by assigning a TF-family to it on the basis of conserved DBDs and integrating the above-mentioned sources of information. Since a protein sequence may have more than one DBD, therefore, for proteins with more than one DBDs, we assigned the TF-family on the basis of DBD with lowest E-value. In order to validate assignment rules and prediction results, we scanned the list of previously known TFs in our classified list of TFs. The known list of TFs was obtained by reviewing the recently published literature.

### Transcription Associated Factor Predictions

For TAF predictions, we performed GO-based analysis of each protein sequence to search for different classes of regulators, i.e. Transcriptional Regulators (TRs), Chromatin Regulators (CRRs) and RNA-regulators (RNARs) among apicomplexans. For TR classification, GO terms with ‘transcription coactivator activity’, ‘transcription corepressor activity’, ‘transcription co-factor activity’ and ‘regulation of transcription’ were used. For CRRs predictions, the GO terms used were ‘chromatin remodeling’, ‘regulation of chromosome’, histone modification ‘histone *ylation’, ‘histone.*ylase activity’ and ‘histone *transferase activity’, as reported by Hong-Mei Zhang et al. [14], whereas, for RNA regulators we used the GO terms-‘RNA-binding’, ‘regulation of transcription by RNA-polymerase’, ‘transcription-RNA dependent’ and ‘transcription initiation from RNA-polymerase’.

### Web interface

The ApicoTFdb web interface has been designed with XHTML, CSS and JavaScript languages. CSS and JavaScript were used for tables and other visualization. In-house PERL scripts perform the database search queries and data retrieval. ApicoTFdb is integrated with a number of additional utilities which facilitates querying the database in more than one way, for instance BLASTn, BLASTp and pdb-BLAST for homology search [34, 35]. Additionally, each of the ApicoTFdb TF entries is dynamically linked to PubMed and Google Scholar for the associated literature search.

## Results

Using the *in-silico* approach, we predicted and classified the TFs and TAFs for 14 apicomplexans species. ApicoTFdb, thus provides a unique platform to analyze several new classes of TFs/TAFs not reported earlier in the parasites genomes.

### TF prediction and classification

#### TF identification using HMM based DBD identification

To predict and classify TFs into their respective superfamily, we identified the conserved DBDs across proteomes of all the target species. In order to achieve the set task, we retrieved all the known DBD profiles from repositories such as AnimalTFDB with 55, DBD database with 147 and FootprintDB with 92 profiles, which were used as reference TF-DBDs profiles for apicomplexans TF identification [S1]. These shortlisted models were then used to scan the DBDs within each of the 14 apicomplexan proteomes. The HMM scanning enabled identification of 1300 putative TFs along with GO based functional analysis, which also resulted in identification of 10 additional putative TFs.

Hence, the above-mentioned TF prediction pipeline and manual curation resulted in the identification of 1310 putative TF proteins representing 22 TF families in the 14 apicomplexans species [Table S1]. The results include, 578 *Plasmodium* spp. TFs, 77 *Cryptosporidium* spp. TFs, overall 648 TFs from *Toxoplasma* spp., *Eimeria* spp., *Cyclospora* spp., *Neurospora* spp., and *Babesia* spp., as shown in Table 1.

#### Comparison with the experimentally verified TFs

All the predicted TFs were manually curated and analyzed for their biological functions. Among the predicted list of TFs, we observed a large number of hypothetical proteins and proteins with unknown functions. Table 1 summarizes the total number of hypothetical/unknown proteins re-annotated as TF in each of the apicomplexan specie. Within our predicted set of TFs, we observed a large number of validated TFs (Table 1). For instance, out of the known 28 *P. falciparum* TFs, we were able to predict 23 annotated TFs. Since we are able to retain majority of known TFs, we extended our analysis to classify this list of TFs according to their respective domains.

#### Genome wide analysis of transcription factors in apicomplexans

Using the TF prediction pipeline, we have successfully assigned functions to 279 proteins, previously annotated as hypothetical, uncharacterized, unspecified product and conserved proteins with unknown function under putative TF class according to their DBDs [Table S2].

Interestingly, the analysis also resulted in the identification TFs of 9 families TUB, NAC, BSD, CCAAT, HTH, Cupin/Jumonji, winged-helix, GO-based and FHA families, not reported earlier for the above apicomplexans (Figure 1). GO based TF-family assignments, integrated with manual curation, enabled inclusion of 10 new TFs, which were missed during our previous conserved DBD domain analysis. As expected, maximum number of TFs (*n*= 401) are from AP2 family, followed by Zn-Finger, General-TF, Myb/SANT, FHA and HMG, with 274, 176, 166, 110 and 64 proteins, respectively, in the 14 apicomplexan species studied here.

**Figure 1:**
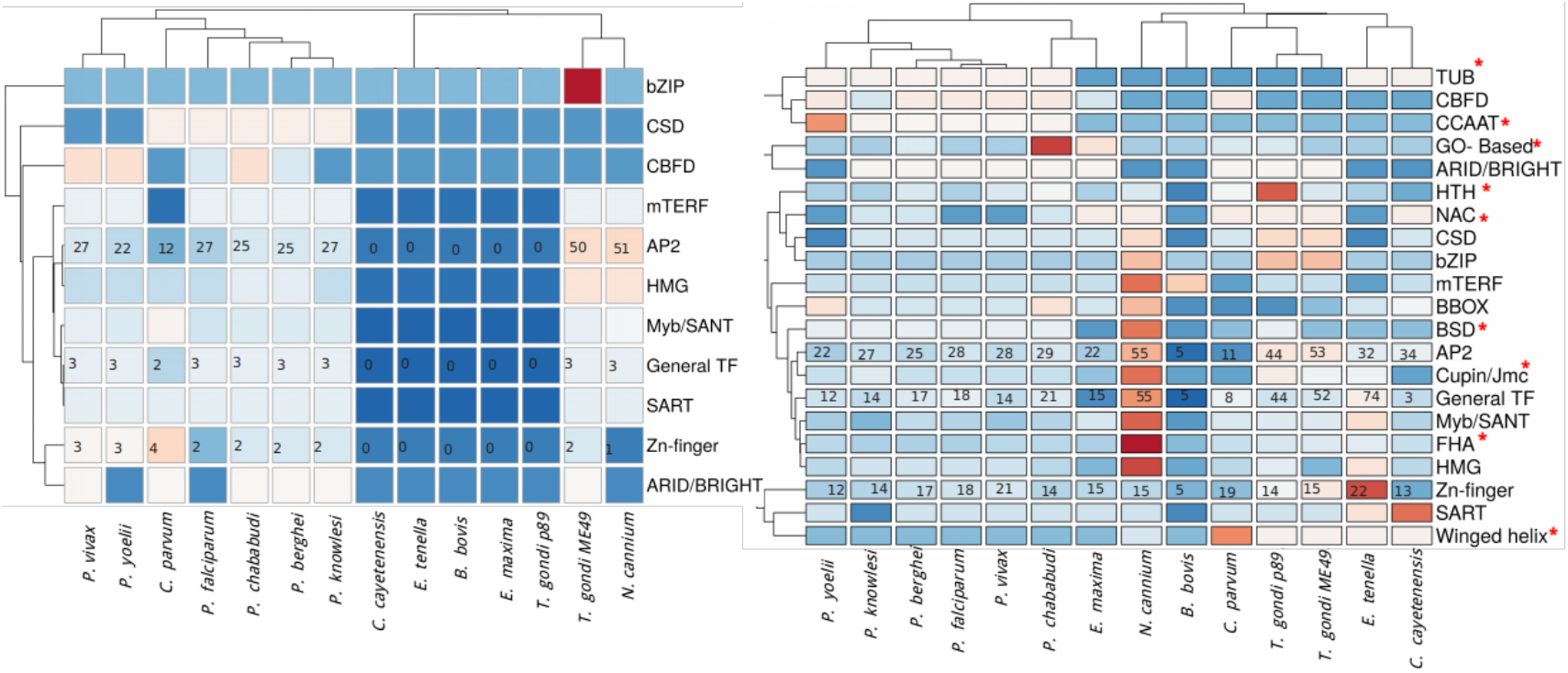
Heat map of TFs in TF families across the apicomplexans, analyzed in the study (a) Distribution of TFs reported earlier in published reports (CisBP, DBD and EupathDB) (b) Distribution of TFs in the current version of ApicoTFdb.

Our analysis also revealed that FHA family is conserved across all the apicomplexans, over-represented in *N. cannium* (with 39 FHA proteins). This family of TFs consists of phosphopeptide-recognition domain and has been identified in eubacterial and eukaryotic genome, unidentified in the archaeal genomes. The FHA family is characterized by its multi-domain architecture in apicomplexans, which includes Zn-Finger C3HC4, Prolyl isomerase (PPIC) and RNA recognition motif (RRM) along with the FHA domain. Apart from TFs, FHA family members include few phosphatases, kinases and RNA-binding proteins, which are involved in many different vital cellular processes [36].

Another family of TF, not reported earlier for apicomplexans, is the TUB. The TUB TFs play important roles in maintaining the functioning of neuronal cells during development and post-differentiation in humans but till date they are not well reported in apicomplexans [37]. Multiple sequence alignment revealed that TUB TF family member harbor conserved Pfam domain PF01167, and have conserved domain architecture in all the species, as shown in Figure 2.

**Figure 2:**
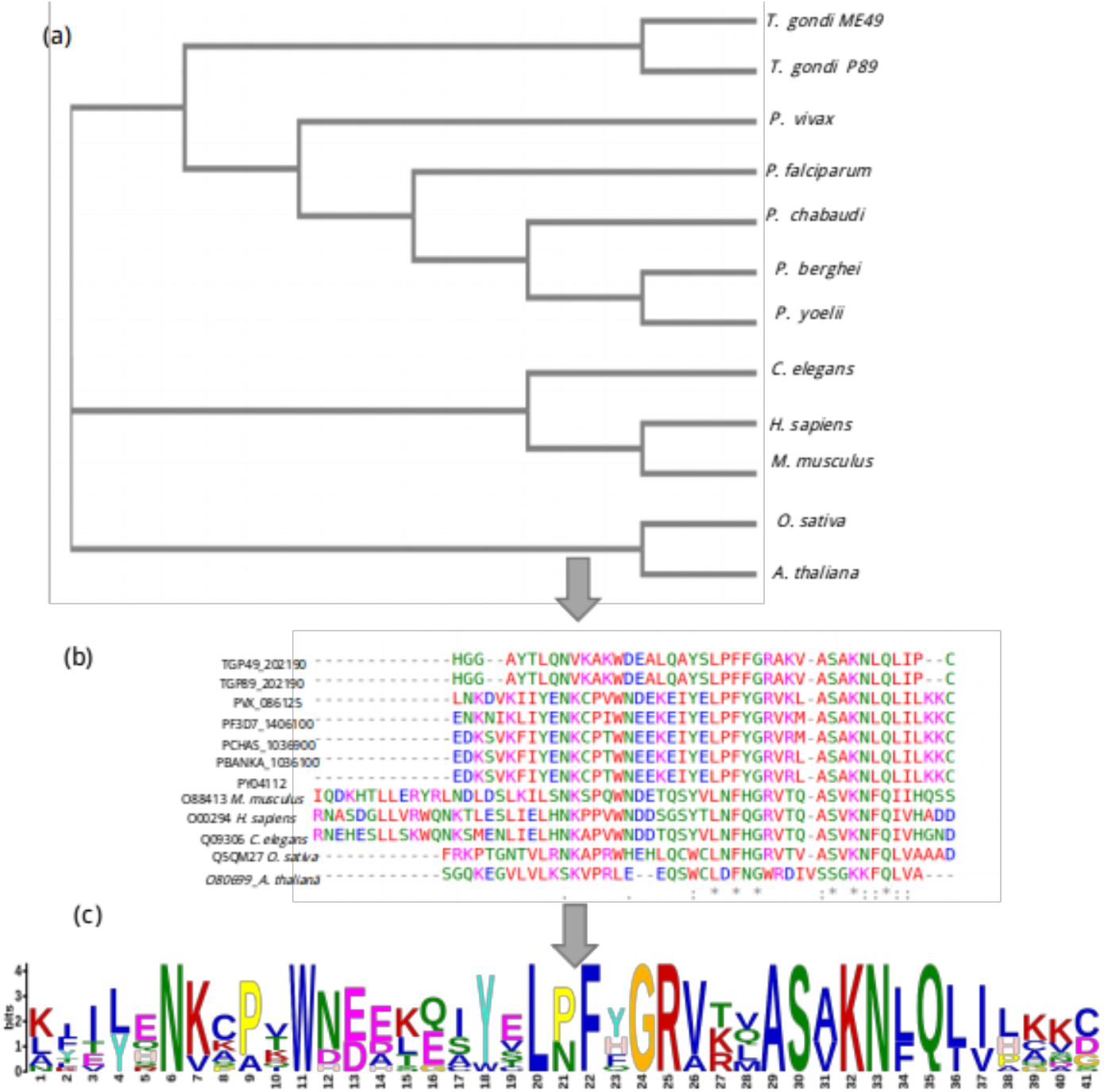
A schematic representation TUB family across 12 species shown here. (a) Phylogenetic analysis of TUB domain showing evolutionary conservation (using NJ method) sharing similar domain architecture (b) Conserved region across TUB domain among different species by multiple sequence alignment (using CLUSTAL-OMEGA) (c) Motif representation of TUB family proteins along with a consensus sequence logo across 12 species.

Intriguingly, we observed that BZIP and winged helix TF family are not present in any of the six Plasmodium species.

### TAF prediction and classification

#### Classification of Transcription-Associated Factors

The TAFs identification pipeline (see materials and methodology) predicted 3047 proteins. In order to remove the false positive predictions (type-I error) across different classes of TAFs [Table S3], manual curation on predicted TAFs was performed. The curation resulted in removal of 797 RNARs proteins with GO functions such as translation, tRNA processing and tRNA modification. Additionally, 125 proteins were found to show GO assigned function for both RNARs and TRs. For further classification, we manually analyzed each protein and classified accordingly in their classes into TRs, CRRs and RNARs (Table 2). This resulted in identification and classification of 666 Transcription Regulators (TRs), 1956 RNARs and 452 chromatin regulators (CRRs), distributed amongst the 14 studied apicomplexan species. Figure 3 summarizes the species-wise distribution of TAFs with most TAF observed in *P knowlesi*. As far as our knowledge, this is a first ever attempt to classify TAFs into TRs, CRRs and RNARs in apicomplexans.

**Table 2:** Manually curated TAFs classification into TRs, RNARs and CRRs.

| **TAFs** | **Initial prediction** | **Manual curation** | **Ortholog Groups** | **Computed GO** | **Curated GO** |
| --- | --- | --- | --- | --- | --- |
| TF-regulators | 678 | 666 | 134 | 124 | 36 |
| RNA-regulators | 2751 | 1956 | 367 | 257 | 87 |
| Chromatin Regulators | 761 | 452 | 113 | 139 | 53 |

**Figure 3:**
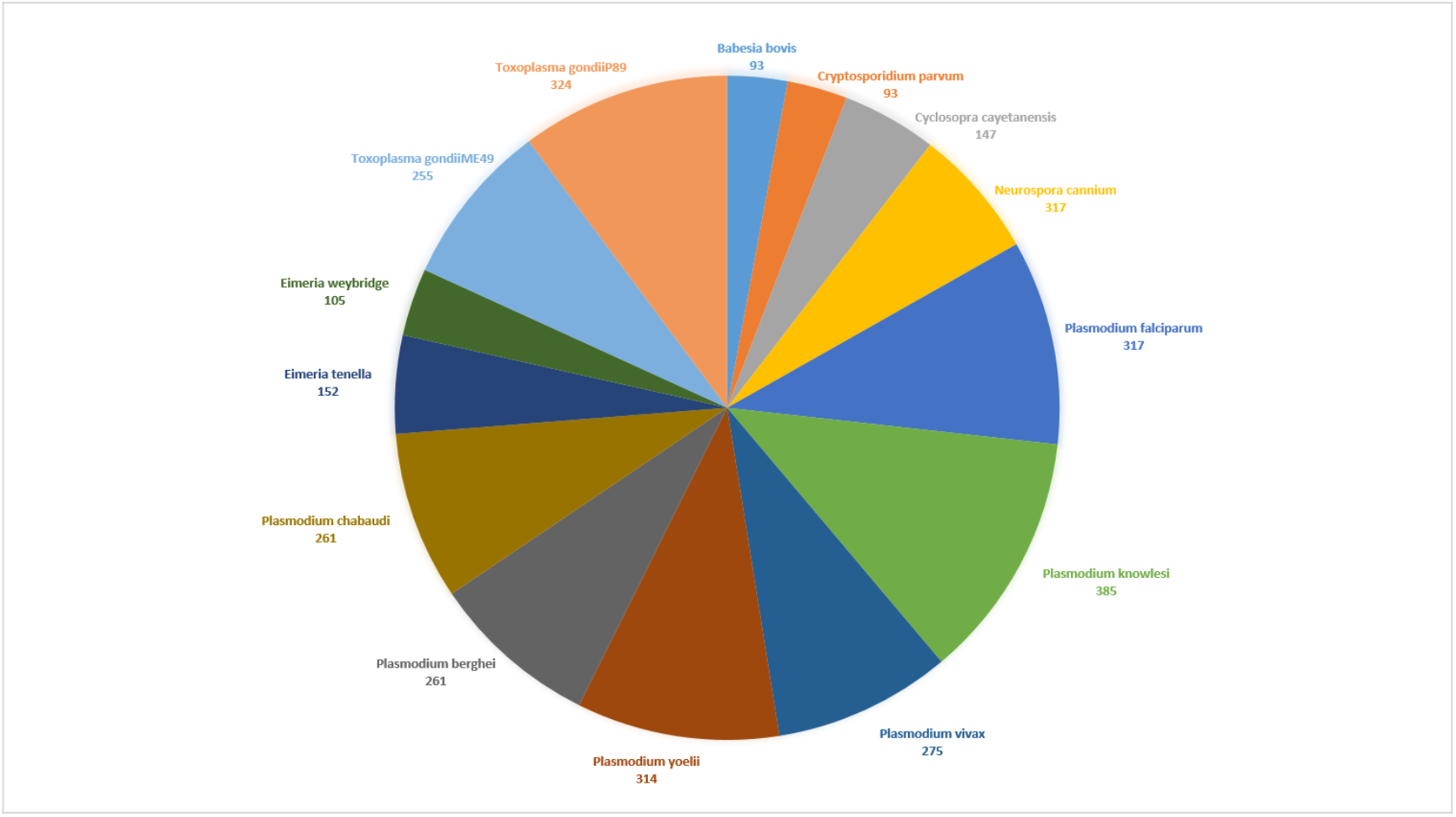
TAFs distribution across apicomplexans

**Figure 4:**
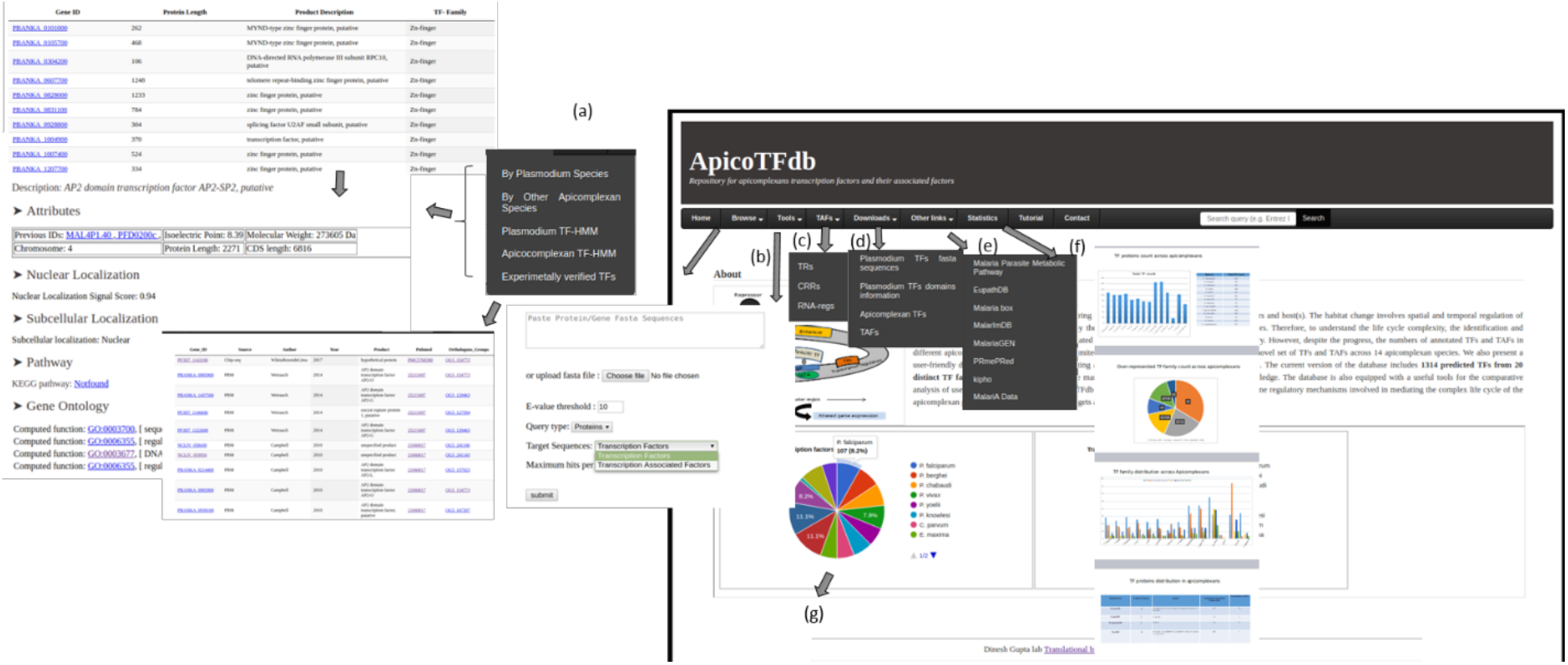
Web interface of ApicoTFdb (a) Browse Section: Browse by species and TF-HMM. By clicking on Gene ID users can retrieve detailed annotation associated with it on gene information page (b) In tool section, users can perform BLAST search using BLASTP and BLASTN against TFs and TAFs (c) TAFs option which is subdivided into TRs, CRRs, and RNARs with their annotation (d) In the downloads section, users can download fasta and tabular annotation file for TFs and TAFs (e) Links option section is linked various other connected external useful resources including KiPho, MPMP, EupathDB, PRmePred, MalariaGen, MalarImDB and Malaria Box (f) Statistics section contains information related to TFs and TAFs count in the current version of ApicoTFdb (g) Dynamic and clickable pie chart showing TFs and TAFs count in ApicoTFdb home page.

#### Web interface and annotations in ApicoTFdb

The ApicoTFdb project is presented as a web-based and user-friendly interface for Apicomplexans TFs and TAFs information retrieval, as shown in Figure 3. The ApicoTFdb database project is organized according to the above described classification of TFs and TAFs into different families based on intrinsic conserved domains and their respective gene ontologies. We included the gene/protein-level information from several relevant web resources including EupathDb and UniProt, Pfam for TF DBD Profile generation. OrthoMCL for orthology profiling; Gene Ontology using AMIGO, PubMed and Google Scholar for related literature information; KEGG (Molecular pathway analysis) and STRING database for protein-protein interactions, which provide necessary information for apicomplexans TFs and TAFs. We have also used CDD [39], PFAM [40], Superfamily [41], and SMART [42] for domain prediction; CELLO2GO for subcellular localization; PATS for apicoplast localization and NCBI-BLAST, including BLASTn, BLASTp and pdb-BLAST, for similarity and homology predictions.

#### Comparison with existing databases and validation

Only a limited number of databases provide information related to the predicted as well as experimentally validated apicomplexan TFs and TAFs, e.g. DBD and CISPB. Though the database is useful, we observed that the information available in CISBP (Database build 1.02) is obsolete as it was implemented via the older version of PlasmoDB (version 10), no longer in use and contains a large set of obsolete IDs. For instance, the database classifies PF14_0010 into the p53 domain, which has been changed to GBP_repeat in the current PlasmoDB annotation (PlasmoDB version 38). Another database viz. DBD with Pfam profile-based prediction also possess limited capabilities and includes only a small set of TFs for the given Apicomplexan species, e.g. only 18 TFs for *P. falciparum*. Table 1 summarizes the number of Apicomplexan TFs reported in ApicoTFdb that compliments the information given in other databases along with the unique set (Table 1). In order to evaluate the confidence of putative transcription factors, we compared our prediction with the published reports. Our findings revealed not only previously predicted and experimentally verified apicomplexans TFs as well as other novel regulatory proteins not reported earlier.

## Discussion and future perspectives

Despite the profound role of transcription regulators in mediating the complex life cycle of apicomplexan parasite across multiple hosts, the number of known and putative regulators in these organisms are exceptionally low. Currently, the repositories like EupathDB are the prime source of putative or known TFs in the parasite genomes. However, most of the available genomes of these parasites are still incomplete and yet to be fully annotated with large number of “hypothetical proteins” and “proteins with unknown function”. Moreover, identification of proteins that can be classified under TFs/TAFs super-family is still a daunting task. Thus, there is an urgent need of a dedicated data resource for retrieving known/putative transcription regulators within different parasite genomes. To identify key transcription regulators in apicomplexan species, we performed an exhaustive scrutinizing of the known DBDs across 14 parasite proteomes. Thereafter, we developed ApicoTFdb, the first exclusive web repository for hosting manually curated TFs and TAFs identified in 14 apicomplexan proteomes.

Amongst the previously identified TFs, the AP2-family TFs are overrepresented in the parasite genomes studied here. Our extended analysis indicates that differences in the distribution of different classes of TFs and TAFs exists across different genomes studies here. We are also successfully able to predict and classify TFs of genomes including *B. bovis, E. maxima, E. tanella, C. cayetanensis and T. gondii ME49* in which extremely limited number of studies till date have done to identify TFs. These species are the causative agent for the intestinal illness in humans, hemorrhagic cecal coccidiosis in young poultry, cattle fever and toxoplasmosis, a worldwide disease that infects one-third of the human population [38].

Moreover, we observed that protein domain information alone is not sufficient to classify a protein under TF or TAFs super-family. Therefore, to restrict the type-I error in our prediction, we classified a given protein under TF or TAF family only after the functional analysis and manual curation, which highlighted several unannotated proteins as TF. For instance, several new TFs and TAFs were identified which were earlier assigned as “Hypothetical proteins” or “Proteins with unknown function”. Thus, the database provides a unique platform to illuminate the list of putative regulators, which otherwise remain obscure with existing portals.

## Acknowledgements

This work was financially supported by the Department of Biotechnology (DBT), Government of India, grants BT/PR6963/BID/7/427/2012 and BT/BI/25/066/2012 awarded to DG.

## Abbreviations

TF: : Transcription Factors
TAF: : Transcription Associated Factors
TR: : Transcription Regulators
CRR: : ChRomatin Regulators
TBP: : TATA box Binding Proteins
DBD: : DNA Binding Domains
PPI: : Protein-Protein Interaction
TUB: : Tubby-family
FHA: : Fork-Head Associated domain

